# Emerging strains of watermelon mosaic virus in Southeastern France: model-based estimation of the dates and places of introduction

**DOI:** 10.1101/2020.10.01.322693

**Authors:** L Roques, C Desbiez, K Berthier, S Soubeyrand, E Walker, E K Klein, J Garnier, B Moury, J Papaïx

## Abstract

Where and when alien organisms are successfully introduced are central questions to elucidate biotic and abiotic conditions favorable to the introduction, establishment and spread of invasive species. We propose a modelling framework to analyze multiple introductions by several invasive genotypes or genetic variants, in competition with a resident population, when observations provide knowledge on the relative proportions of each variant at some dates and places. This framework is based on a mechanistic-statistical model coupling a reaction-diffusion model with a probabilistic observation model. We apply it to a spatio-temporal dataset reporting the relative proportions of five genetic variants of watermelon mosaic virus (WMV, genus *Potyvirus*, family *Potyviridae*) in infections of commercial cucurbit fields. Despite the parsimonious nature of the model, it succeeds in fitting the data well and provides an estimation of the dates and places of successful introduction of each emerging variant as well as a reconstruction of the dynamics of each variant since its introduction.

## INTRODUCTION

Plant and animal species – and as a consequence, their pathogens and pests – are translocated across the globe at an ever-increasing rate since the 19th century [1]. Most introductions of microorganisms and insects (notably plant pathogens and/or their arthropod vectors) are accidental, while a few are deliberate for biological control purposes [2]. Introductions of plant pathogens can have a dramatic impact on agricultural production and food security (Anderson et al., 2004) and being able to predict the fate of a newly introduced disease based on surveillance data is pivotal to anticipate control actions [3].

Where and when alien organisms are successfully introduced are central questions for the study of biological invasions, to elucidate biotic and abiotic conditions favorable to the introduction and establishment and spread of invasive species [4, 5]. Indeed, unraveling such conditions is a prerequisite to map the risk of invasion and to design an efficient surveillance strategy [5] as initial phases are critical for the establishment success of introduced pathogens [6, 7]. It is also during this phase that invaders are the easiest to control [4, 8]. In addition, a precise knowledge of the dates and places of introduction is critical to accurately determine invader reproduction and dispersal parameters, as their estimated values are directly dependent on the time and distance between the actual introduction and the observations [9, 10]. However, biological invasions by alien organisms are often reported several years after the initial successful introduction event [11]. Thus, monitoring data generally do not provide a clear picture of the date and place of introduction.

Many successful emergences of plant viruses have taken place worldwide in the last decades [12, 13], whereas some have only been observed punctually with no long term establishment of the pathogens [14]. As plant viruses are rapidly-evolving pathogens with small genomes and high mutation rates [15], they may present measurably evolving populations [16] over the time scale of decades. Thus, the spatio-temporal histories of invading species can sometimes be reconstructed from georeferenced and dated genomic data, with phylogeographic methodologies [17, 18, 19] or population dynamic models embedding evolutionary processes. Over shorter time-scales, or when only abundance data are available, recent mechanistic modelling approaches have been proposed to infer the date and place of introduction of a single species along with other demographic parameters [20]. In many modeling approaches, the interactions of the new viruses or new strains with preexisting virus populations are not taken into consideration, even if it is known that synergism or competition between virus species or strains can affect their maintenance and spread [21, 22, 23].

Mechanistic models are increasingly used in statistical ecology because, compared to purely correlative approaches, their parameters can directly inform on biological processes and life history traits. Among them, the reaction-diffusion framework is widely used in spatial ecology to model dynamic species distributions [24, 25]. In such models, the diffusion coefficient is related to dispersal ability and the growth or competition coefficients may help in understanding the respective interactions between different species, variants or genotypes. Additionally, reaction-diffusion models can easily account for spatio-temporal heterogeneities [26, 27]. A drawback of this type of approach is that the compartments that are modelled typically correspond to continuous population densities, which rarely match with observation data. There is therefore a challenge in connecting the solution of the model with complex data, such as noisy data, binary data, temporally and spatially censored data. Recent approaches have been proposed to bridge the gap between reaction-diffusion models and data (e.g., [28], in a framework known as mechanistic-statistical modelling [29, 30, 31]. An advantage of this method is that it allows to estimate simultaneously both the biological parameters and the date and place of introduction in a single framework [20]. Recently, this type of approach was applied to localize and date the invasion of South Corsica by *Xylella fastidiosa*, based on a single-species reaction-diffusion model and binary data [9].

Our objective here is to propose a modelling framework to deal with multiple introductions by several invasive variants, in competition with a resident population, when observations provide knowledge on the relative proportions of each variant at some dates and places rather than absolute abundances of the different variants. We develop a reaction-diffusion mechanistic-statistical model applied to a genetic spatio-temporal dataset reporting the relative proportions of five genetic variants of watermelon mosaic virus (WMV, genus *Potyvirus*, family *Potyviridae*) in infections of commercial cucurbit fields. Our framework allows us to (i) estimate the dates and places of successful introduction of each emerging variant along with other ecological parameters, (ii) reconstruct the invasion history of the emerging variants from their introduction sites, (iii) detect competitive advantages of the emerging variants as compared to the resident population, and (iv) predict the fate of the different genetic groups, in particular the takeover of the emerging variants over the resident population.

## MATERIAL AND METHODS

### Data

#### Pathosystem

WMV is widespread in cucurbit crops, mostly in temperate and Mediterranean climatic regions of the world [14]. WMV has a wide host range including some legumes, orchids and many weeds that can be alternative hosts [14]. Like other potyviruses, it is non-persistently transmitted by at least 30 aphid species [14]. In temperate regions, WMV causes summer epidemics on cucurbit crops, and it can overwinter in several common non-cucurbit weeds when no crops are present [14, 32]. WMV has been common in France for more than 40 years, causing mosaics on leaves and fruits in melon, but mostly mild symptoms on zucchini squash. Since 2000, new symptoms were observed in southeastern France on zucchini squash: leaf deformations and mosaics, as well as fruit discoloration and deformations that made them unmarketable. This new agronomic problem was correlated to the introduction of new molecular groups of WMV strains. At least four new groups have emerged since 2000 and they have rapidly replaced the native “classical” strains, causing important problems for the producers [33].

In this study, we focus on the pathosystem corresponding to a classical strain (CS) and four emerging strains (ES_k_, *k* = 1, …, 4) of WMV and their cucurbit hosts.

#### Study area and sampling

The study area, located in Southeastern France, is included in a rectangle of about 25000 km^2^ and is bounded on the South by the Mediterranean Sea. Between 2004 and 2008, the presence of WMV had been monitored in collaboration with farmers, farm advisers and seed companies. Each year, cultivated host plants were collected in different fields and at different dates between May 1^st^ and September 30^th^. In total, more than two thousand plant samples were collected over the entire study area. All plant samples were analyzed in the INRAE Plant Pathology Unit to confirm the presence of WMV and determine the molecular type of the virus strain causing the infection (see [33] for detail on field and laboratory protocols). All infected host plants were cucurbits, mostly melon and different squashes (e.g., zucchini, pumpkins).

#### Observations

In the absence of individual geographic coordinates, all infected host plants were attributed to the centroid of the municipality (French administrative unit, median size about 10 km^2^) where they have been collected. Then for one date, one observation corresponded to a municipality in which at least one infected host plant was sampled. Table 1 summarizes for each year, the number of observations (i.e. number of municipalities), the number of infected plants sampled and the proportion of each WMV strain (CS, and ES_1_ to ES_4_) found in the infected host plants. Errors in assignment of virus samples to the CS or ES strains was negligible because of the large genetic distance separating them: 5 to 10 % nucleotide divergence both in the fragment used in the study and in complete genomes [33], also precluding the possibility of local jumps between groups by accumulation of mutations.

**Table 1.** Number of observations and corresponding proportions of classical and emerging strains.

|  | 2004 | 2005 | 2006 | 2007 | 2008 |
| --- | --- | --- | --- | --- | --- |
| # observations | 67 | 64 | 68 | 50 | 40 |
| # infected samples | 408 | 371 | 422 | 280 | 212 |
| Classical strain | 55% | 45% | 28% | 17% | 14% |
| Emerging strain 1 | 21% | 23% | 22% | 37% | 27% |
| Emerging strain 2 | 13% | 18% | 23% | 21% | 32% |
| Emerging strain 3 | 1% | 4% | 3% | 5% | 3% |
| Emerging strain 4 | 10% | 10% | 24% | 20% | 24% |

#### Landscape

To approximate the density of WMV host plants over the study area, we used 2006 land use data (i.e. BD Ocsol 2006 PACA and LR) produced by the CRIGE PACA (http://www.crige-paca.org/) and the Association SIG-LR (http://www.siglr.org/lassociation/la-structure.html). Based on satellite images, land use is determined at a spatial resolution of 1/50,000 using an improved three-level hierarchical typology derived from the European Corine Land Cover nomenclature. Here we used the third hierarchical level of the BD Ocsol typology (i.e. 42 land use classes) to classify the entire study area in three habitats: 1) WMV-susceptible crops, 2) habitats unfavorable to WMV host plants (e.g. forests, industrial and commercial units…) and, 3) non-terrestrial habitat (i.e. water). The proportion of WMV-susceptible crops was then computed within all cells of a raster covering the entire study area, with a spatial resolution of 1.4 × 1.4 km^2^. These proportions were used to approximate host plant density *z*(***x***), which was normalized, so that *z*(***x***) = 0 corresponds to an absence of host plants and *z*(***x***) = 1 to the maximum density of host plants (**Fig. 1**).

**Fig. 1.**
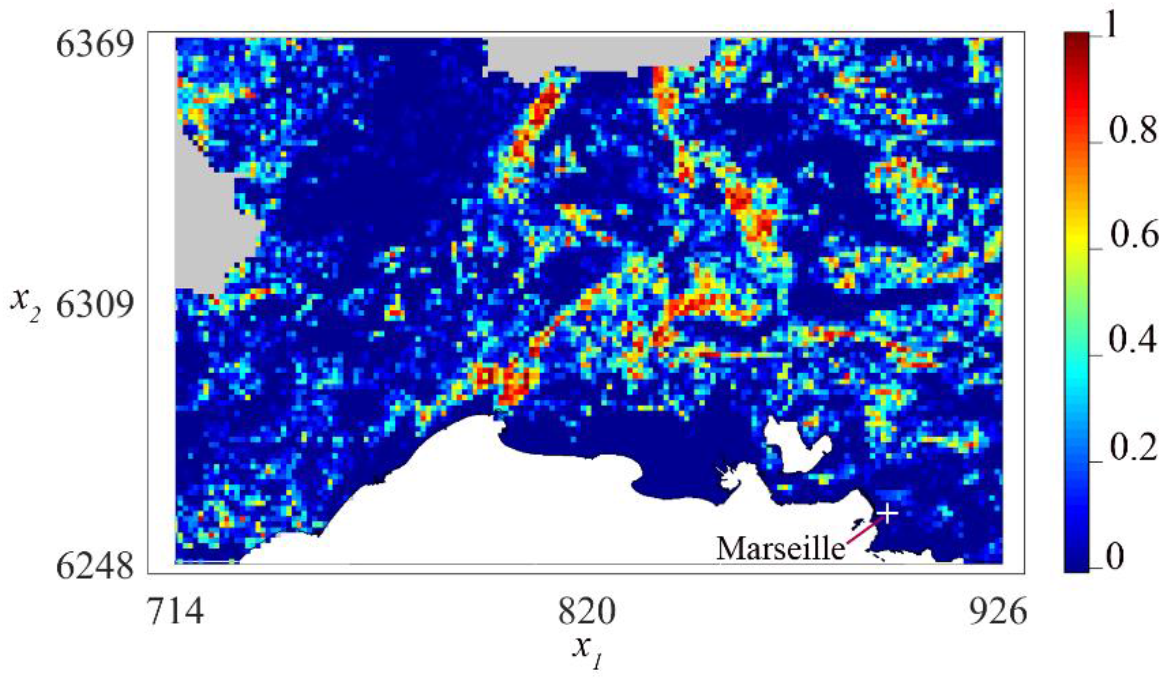
Approximated density *z*(***x***) of the host plants in the study area. The density was normalized, so that *z*(***x***) = *z*(*x*_1_, *x*_2_) = 0 corresponds to an absence of cucurbit plants and *z*(***x***) = 1 to the maximum density. The axes *x*_1_ and *x*_2_ correspond to Lambert93 coordinates (in km). The white regions are non-terrestrial habitats (water). Land use data were not available in the gray regions; the host plant density was then computed by interpolation.

### Mechanistic-statistical model

The general modeling strategy is based on a mechanistic-statistical approach [31, 20, 10]. This type of approach combines a mechanistic model describing the dynamics under investigation with a probabilistic model conditional on the dynamics, describing how the measurements have been collected. This method that has already proved its theoretical effectiveness in determining dispersal parameters using simulated genetic data [10] aims at bridging the gap between the data and the model for the determination of virus dynamics.

Here, the mechanistic part of the model describes the spatio-temporal dynamics of the virus strains, given the model parameters (demographic parameters, introduction dates/sites). This allows us to compute the expected proportions of the five types of virus strains (CS and ES_1_ to ES_4_) at each date and site of observation. The probabilistic part of the mechanistic-statistical model describes the conditional distribution of the observed proportions of the virus strains, given the expected proportions. Using this approach, it is then possible to derive a numerically tractable formula for the likelihood function associated with the model parameters.

### Population dynamics

The model is segmented into two stages: (1) the intra-annual stage describes the dispersal and growth of the five virus strains during the summer epidemics on cucurbit crops, and the competition between them, during a period ranging from May 1^st^ (noted *t* = 0) to September 30 (noted *t* = *t*_*f*_, *t*_*f*_ = 153 days); (2) the inter-annual stage describes the winter decay of the different strains when no crops are present and the virus overwinters in weeds. We denote by *c*^*n*^(*t*, ***x***) and 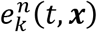 the densities of classical strain (CS) and emerging strains (ES_k_, *k* = 1, …, 4), at position ***x*** and at time *t* of year *n*.

#### Dynamics of the classical strain before the first introduction events

Before the introduction of the first emerging strain, the intra-annual dynamics of the population CS is described by a standard diffusion model with logistic growth: *∂*_*t*_*c*^*n*^ = *D*Δ*c*^*n*^ + *rc*^*n*^(*z*(***x***) − *c*^*n*^). Here, Δ is the Laplace 2D diffusion operator (sum of the second derivatives with respect to coordinate). This operator describes uncorrelated random walk movements of the viruses, with the coefficient *D* measuring the mobility of the viruses (e.g., [24, 34]). The term *r z*(***x***) is the intrinsic growth rate (i.e., growth rate in the absence of competition) of the population, which depends on the density of host plants *z*(***x***) and on a coefficient *r* (intrinsic growth rate at maximum host density). Under these assumptions, the carrying capacity at a position ***x*** is equal to *z*(***x***), which means that the population densities are expressed in units of the maximum host population density. The model is initialized by setting *c*^1980^(0, ***x***) = (1 − *m*_*c*_) *z*(***x***), where *m*_*c*_ is the winter decay rate of the CS (see the description of the inter-annual stage below). In other terms, we assume that the CS density is at the carrying capacity in 1979, i.e., 5 years after its first detection and 20 years before the first detections of ESs [35].

#### Introduction events

The ESs are introduced during years noted *n*_*k*_ ≥ 1981, at the beginning of the intra-annual stage (other dates of introduction within the intra-annual stage would lead – at most – to a one-year lag in the dynamics). Their densities are 0 before introduction: 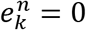 for *n* < *n*_*k*_. Once introduced, the initial density of any ES is assumed to be 1/10^th^ of the carrying capacity at the introduction point (other values have been tested without much effect, see Supplementary Fig. S1), with a decreasing density as the distance from this point increases:

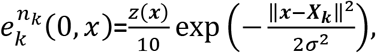

where ***X*_*k*_** is the location of introduction of the strain *k*. In our computations, we took σ = 5 km for the standard deviation.

#### Intra-annual dynamics after the first introduction event

Intra-annual dynamics were described by a neutral competition model with diffusion (properties of this model have been analyzed in [RGHK12]):

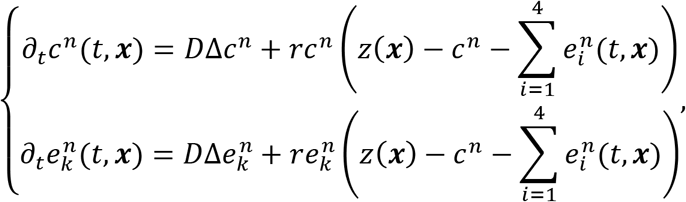

for *t* = 0 … *t*_*f*_ and for all introduced emerging strains, i.e. all *k* such that *n* ≥ *n*_*k*_. We assume reflecting boundary conditions, meaning that the population flows vanish at the boundary of the study area, due to truly reflecting boundaries (e.g., sea coast in the southern part of the site) or symmetric inward and outward fluxes [24]. In addition, in order to limit the number of unknown parameters, and in the absence of precise knowledge on the differences between the strains, we assume here that the diffusion, competition and growth coefficients are common to all the strains during the intra-annual stage (see the discussion for more details on this assumption).

#### Inter-annual dynamics

The population densities at time *t* = 0 of year *n* are connected with those of year *n* − 1, at time *t* = *t*_*f*_, through the following formulas:

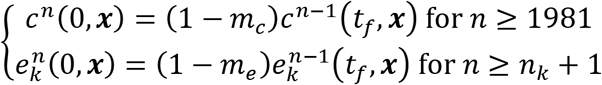

with *m*_*c*_ and *m*_*e*_ the winter decay rates of the CS and ESs strains (note that *m*_*e*_ is common to all of the ESs). Estimation of CS and ES decay rates provides an assessment of the competitive advantage of one type of strain vs the other.

#### Numerical computations

The intra-annual dynamics were solved using Comsol Multiphysics^®^ time-dependent solver, which is based on a finite element method (FEM). The triangular mesh which was used for our computations is available as Supplementary Fig. S2.

### Probabilistic model for the observation process

During the years *n* = 2004, …, 2008, *I*_*n*_ observations were made (see Section *Observations* above and **Table 1**). They consist in counting data, that we denote by *C*_*i*_ and *E*_*k,i*_ for *k* = 1, … ,4 and *i* = 1, … , *I*_*n*_, corresponding to the number of samples infected by the CS and ESs strains, respectively, at each date of observation and location (*t*_*i*_, ***x***_*i*_). Note that these dates and locations depend on the year of observation *n*. More generally, the above quantities should be noted 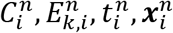for simplicity, the index *n* is omitted in the sequel, unless necessary.

We denote by 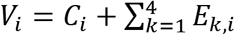 the total number of infected samples observed at (*t*_*i*_, ***x***_*i*_). The conditional distribution of the vector (*C*_*i*_, *E*_1,*i*_, *E*_2,*i*_, *E*_3,*i*_, *E*_4,*i*_), given *V*_*i*_ can be described by a multinomial distribution 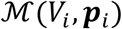 with 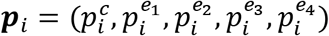 the vector of the respective proportions of each strain in the population at (*t*_*i*_, ***x***_*i*_). We chose to work conditionally to *V*_*i*_ because the sample sizes are not related to the density of WMV.

### Statistical inference

#### Unknown parameters

We denote by Θ the vector of unknown parameters: the diffusion coefficient *D*, the intrinsic growth rate at maximum host density *r*, the winter decay rates (*m*_*c*_, *m*_*e*_) and the locations (*x*_*k*_ ∈ ℝ^2^) and years (*n*_*k*_) of introduction, for *k* = 1, …, 4. Thus Θ ∈ ℝ^16^.

#### Computation of a likelihood

Given the set of parameters Θ, the densities *c*^*n*^(*t*, ***x***|Θ) and 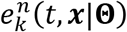 can be computed explicitly with the mechanistic model for population dynamics. Thus, at a given year *n*, at (*t*_*i*_, *x*_*i*_), the parameter *p*_*i*_ of the multinomial distribution 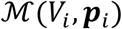 writes:

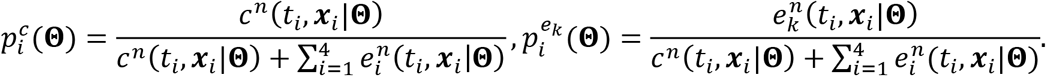

The probability *P*(*C*_*i*_, *E*_1,*i*_, *E*_2,*i*_, *E*_3,*i*_, *E*_4,*i*_|Θ, V_i_) of the observed outcome *C*_*i*_, *E*_1,*i*_, *E*_2,*i*_, *E*_3,*i*_, *E*_4,*i*_ is then

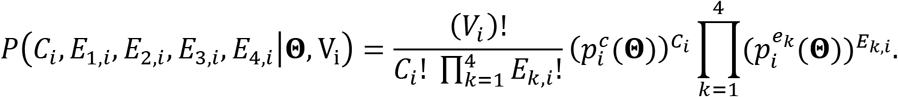

Assuming that the observations during each year and at each date/location are independent from each other conditionally on the virus strain proportions, we get the following formula for the likelihood:

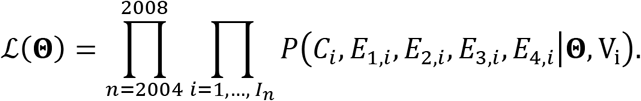

#### A priori constraints on the parameters

By definition and for biological reasons, the parameter vector Θ satisfies some constraints. First, *D* ∈ (10^−4^, 10) km^2^/day, *r* ∈ (0.1,1) day^−1^, and *m*_*c*_, *m*_*e*_ ∈ {0,0.1,0.2, …, 0.9}, (see Supplementary Note S7 for a biological interpretation of these values). Second, we assumed that the locations of introductions ***X*_*k*_** belong to the study area. To facilitate the estimation procedure, the points ***X*_*k*_** were searched in a regular grid with 20 points (see Supplementary Fig. S3), and the dates of introduction *n*_*k*_ were searched in {1985,1990,1995,2000}.

#### Inference procedure

Due to the important computation time (4 minutes in average for one simulation of the model on an Intel(R) Core(R) CPU i7-4790 @ 3.60GHz), we were not able to compute an *a posteriori* distribution of the parameters in a Bayesian framework. Rather, we used a simulated annealing algorithm to compute the maximum likelihood estimate (MLE), i.e., the parameter Θ^*^ which leads to the highest log-likelihood. This is an iterative algorithm, which constructs a sequence (Θ_*j*_)_*j*≥1_ converging in probability towards a MLE. It is based on an acceptance-rejection procedure, where the acceptance rate depends on the current iteration *j* through a “cooling rate” (*α*). Empirically, a good trade-off between quality of optimization and time required for computation (number of iterations) is obtained with exponential cooling rates of the type *T*_0_ *α*^*j*^ with 0 < *α* < 1 and some constant *T*_0_ ≫ 1 (this cooling schedule was first proposed in =[36]=[36]). Too rapid cooling (*α* ≪ 1) results in a system frozen into a state far from the optimal one, whereas too slow cooling (*α* ≈ 1) leads to important computation times due to very slow convergence. Here, we ran 6 parallel sequences with cooling rates *α* ∈ {0.995,0.999,0.9995}. For this type of algorithm, there are no general rules for the choice of the stopping criterion [HenJac03], which should be heuristically adapted to the considered optimization problem. Here, our stopping criterion was that Θ_*j*_ remained unchanged during 500 iterations. The computations took about 100 days (CPU time).

#### Confidence intervals and goodness-of-fit

To assess the model’s goodness-of-fit, 95% confidence regions were computed for the observations (*C*_*i*_, *E*_1,*i*_, *E*_2,*i*_, *E*_3,*i*_, *E*_4,*i*_) at each date/location (*t*_*i*_, ***x***_*i*_), and for each year of observation. The confidence regions were computed by assessing the probability of each possible outcome of the observation process, at each date/location, based on the computed proportions 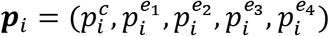, corresponding to the output of the mechanistic model using the MLE Θ^*^ and given the total number of infected samples *V*_*i*_. Then, we checked if the observations belonged to the 95% most probable outcomes.

## RESULTS

### Convergence and goodness-of-fit

As expected, the highest likelihood was obtained with a slow cooling rate (*α* = 0.9995). The corresponding MLE, denoted by 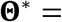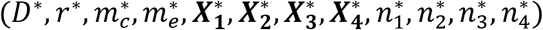 is presented in **Table 2**. In average, 96% of the observations fell within the 95% confidence regions, indicating that the model fits the data well (277 observations over a total of 289; 94% in 2004, 98% in 2005, 96% in 2006, 96% in 2007 and 95% in 2008). We also note that the likelihood function is peaked at Θ^*^ in the sense that any perturbation in one component of Θ^*^ leads to lower likelihood (see Supplementary Note S8 for likelihood-ratio based confidence intervals and Supplementary Fig. S4 for more details on the profile of the likelihood function). This suggests that the MLE Θ^*^ is close to the actual maximizer of the likelihood function.

**Table 2.** Maximum likelihood estimates.

| Biological parameter | $D^*$ | $r^*$ | $m_c^*$ | $m_e^*$ |
| --- | --- | --- | --- | --- |
| Value | $0.44 \text{ km}^2 \text{ day}^{-1}$ | $0.31 \text{ day}^{-1}$ | $0.5 \text{ year}^{-1}$ | $0 \text{ year}^{-1}$ |
| Date of introduction | $n_1^* (ES1)$ | $n_2^* (ES2)$ | $n_3^* (ES3)$ | $n_4^* (ES4)$ |
| Value | 1990 | 1990 | 1990 | 1995 |
| Site of introduction | $X_1^*$ | $X_2^*$ | $X_3^*$ | $X_4^*$ |
| Value (Lambert 93, km) | (926,6369) | (926,6369) | (758,6369) | (758,6294) |

### Parameter values

As shown in **Table 2**, the MLE corresponds to the same date of introduction for ES_1_, ES_2_, ES_3_ whereas ES_4_ has been introduced five years later. Note that a same date of introduction does not mean a same date of detection: depending on local conditions, some strains may establish and spread faster than others (as observed below for ES_4_).

Regarding the sites of introduction, the MLE indicates that ES_1_ and ES_2_ have been introduced at the northeastern corner of the study area, ES_3_ in the northwest and ES_4_ in the southwest (see the white crosses in the 2004 panel of **Fig. 2**; see also Supplementary Fig. S3 for more details on the corresponding likelihood). Three introduction points (ESs 1, 2 and 3) were estimated at the edge of the study site, indicating that the introductions may have occurred outside of the study area.

**Fig. 2.**
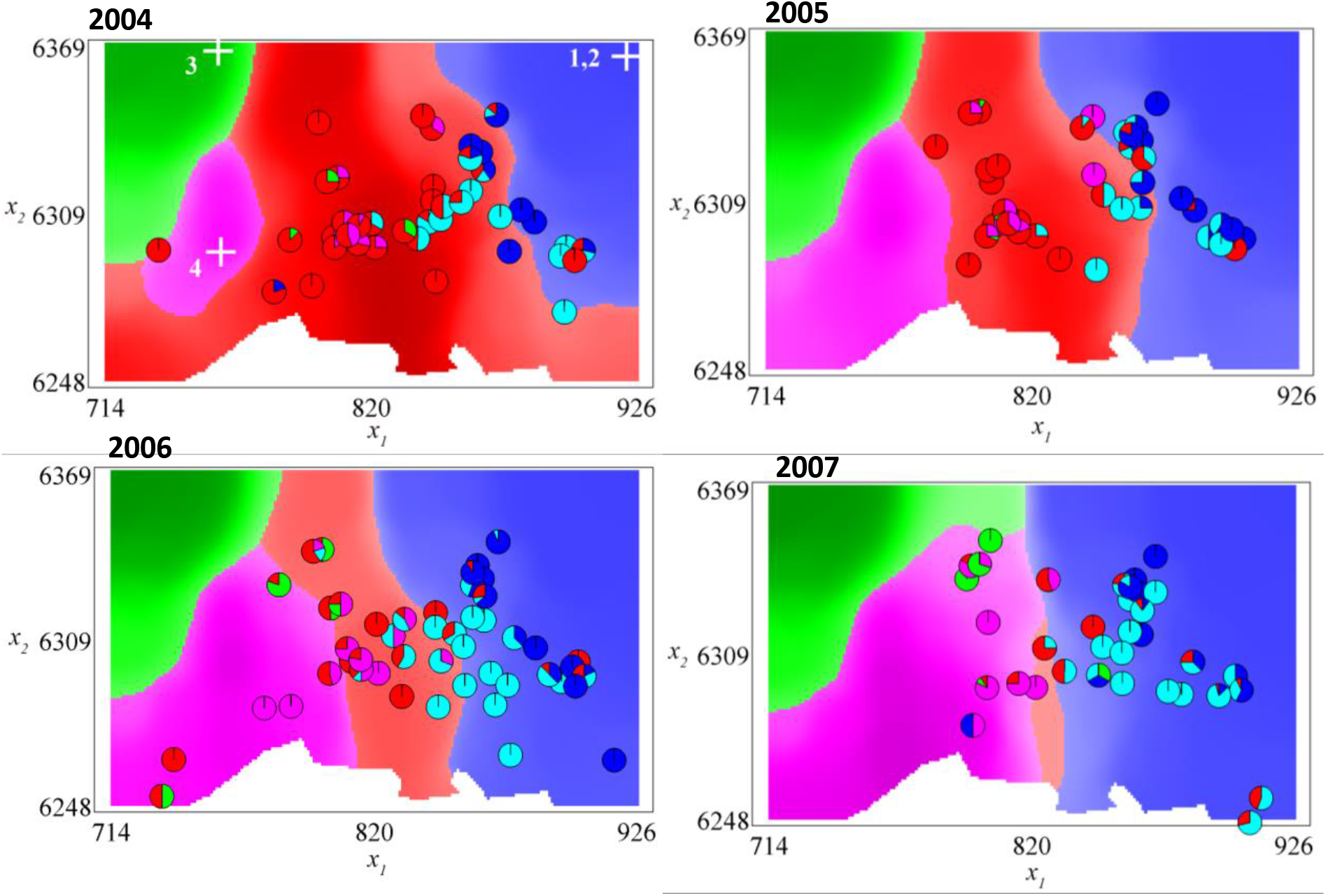
Proportions of the classical and emerging strains in the landscape: data and simulations. The colors of the shaded regions indicate which strain is *the most prevalent*. The red regions correspond to the CS strain; light blue and blue: ES_1_, ES_2_ (these two strains have the same density, only ES_1_ is represented); green: ES_3_; pink: ES_4_. The pie charts describe the relative proportions of the strains found in the data (same color legend). The white crosses on the 2004 panel represent the estimated sites of introduction. The simulation results presented here correspond to the middle of the intra-annual stage (2^nd^ week of June), and were obtained with the MLE Θ^*^.

Regarding the biological parameters, the winter decay rate of the CS strain was much higher than that of the ESs strains (0.5 vs. 0), reflecting a high competitive advantage of the ESs. The value *D*^*^ = 0.44 km^2^ per day of the diffusion coefficient indicates that a virus travels about 1.2 km per day in average during the growing season of cucurbit crops (see Supplementary Note S7). The growth rate of 0.31 day^−1^ corresponds to an increase by a factor *e*^0.31^ ≈ 1.3 each day, in the absence of competition.

### Strain distributions

**Fig. 2** depicts the most prevalent strains at each position in the landscape, during four of the five years of observation (2008 is presented in Supplementary Fig. S5), obtained by solving our model with the MLE Θ^*^, together with the data. We graphically note a good agreement between the positions of the observed strains and the distributions obtained with the model: ES_1_ and ES_2_ tend to be distributed in the eastern part of the study area, ES_3_ in the northwestern part and ES_4_ in the southwestern part, while the CS strain tends to be progressively confined to the central part of the study area. In 2007, ES_3_ seems to be more prevalent according to the model than suggested by the observations, probably due to its introduction site, which is far from the observation sites (see the first panel in **Fig. 2**). Note that, with the deterministic framework used here, as ES_1_ and ES_2_ share the same date and position of introduction, and the same parameter values, their distributions are completely equal; thus only ES_1_ is represented in the Figures.

The spatial distributions of the different strains at each year where one of the emerging strains becomes locally more prevalent are depicted in **Fig. 3**. Although ESs 1, 2 and 3 have been introduced at the same date, their dynamics are influenced by local conditions: ESs 1 and 2 become the most prevalent at least in one part of the study area in 1996 (6 years after their introduction), ES_3_ in 2001 (11 years after introduction) and ES_4_ in 2002 (7 years after its introduction). Thus, despite the neutrality assumption, the heterogeneity of the landscape leads to different durations of the establishment stage. The full timeline of the dynamics of the different strain proportions in the landscape, from the first estimated introduction date of an emerging strain (1990) to 2019 is available as Supplementary Fig S6. Since 2008, due to their competitive advantage (modelled here as a lower winter decay rate), the ESs replaced the CS, which is not anymore the most prevalent strain, whatever the position in the study area, 18 years after the first introduction. Before saturation, the spread rates of the ESs are about 5km/year (estimated as the slope of 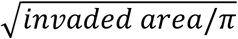, the invaded area corresponding to virus densities> carrying capacity/100). Then, the distribution of the ESs remains almost at equilibrium until the last year of simulation, which is a consequence of the neutrality assumption (equal fitness of all the ESs) (**Fig. 3**; last panel).

**Fig. 3.**
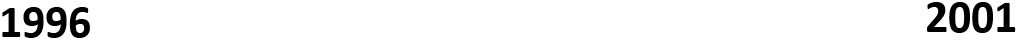
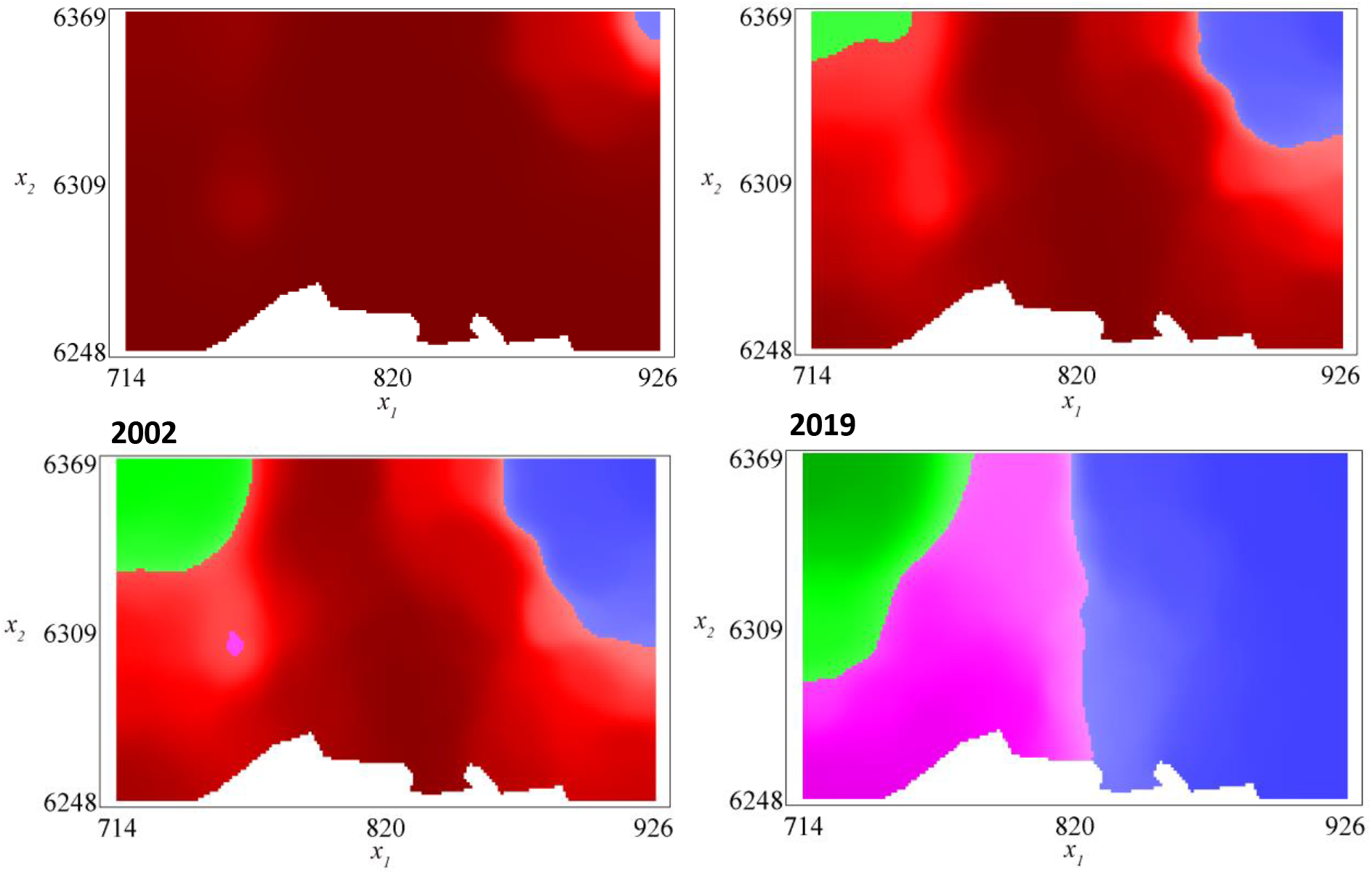
Simulated proportions of the classical and emerging strains in the landscape: before and after the observation window. The simulation results presented here correspond to the middle of the intra-annual stage (2^nd^ week of June), and were obtained with the MLE Θ^*^. The colors of the shaded regions indicate which strain is the most prevalent. The red regions correspond to the classical strain; blue: ES_1_, ES_2_ (these two strains have the same density, only ES_1_ is represented); green: ES_3_; pink: ES_4_.

#### Average proportions in the study area and effect of the CS on the ESs

To get a quantitative insight into the replacement of the CS by the ESs, we computed the relative global proportion of each strain by integrating the simulation results (with the parameters corresponding to the MLE Θ^*^) over the study area (**Fig. 4**, panel (A)). Before the first introduction in 1990, the classical strain represents 100% of the infections. In 2010, it represents only 10% of the infections. This decline, which was already visible in the 2004-2008 data [33, 32], is well-captured by the model, though with a slight advance. These discrepancies between the predicted proportions and the data are probably due to the positions of the observation sites, which are concentrated at the center of the domain, where the CS is more prevalent (see **Fig. 2**). In order to understand the effect of the presence of a resident CS strain on the emerging ESs strains, we compared the dynamics presented in **Fig. 4**, panel (A) with a hypothetical scenario describing the dynamics of the ESs in the absence of CS. For this, we used the MLE Θ^*^, to simulate the hypothetical dynamics of the ESs assuming that the CS density is 0. The results are depicted in **Fig. 4**, panel (B). We observe a very fast convergence to an equilibrium, compared to the situation where the CS is present. Additionally, the last introduced ES (ES_4_) cannot establish, and ES_3_ which was confined in an unfavorable region in the presence of the CS, reaches more favorable regions, leading to a higher proportion. Thus, the competition with the CS alters the outcome of the competition between the ESs, and seems to promote the diversity of the ESs by slowing down the overall dynamics.

**Fig. 4.**
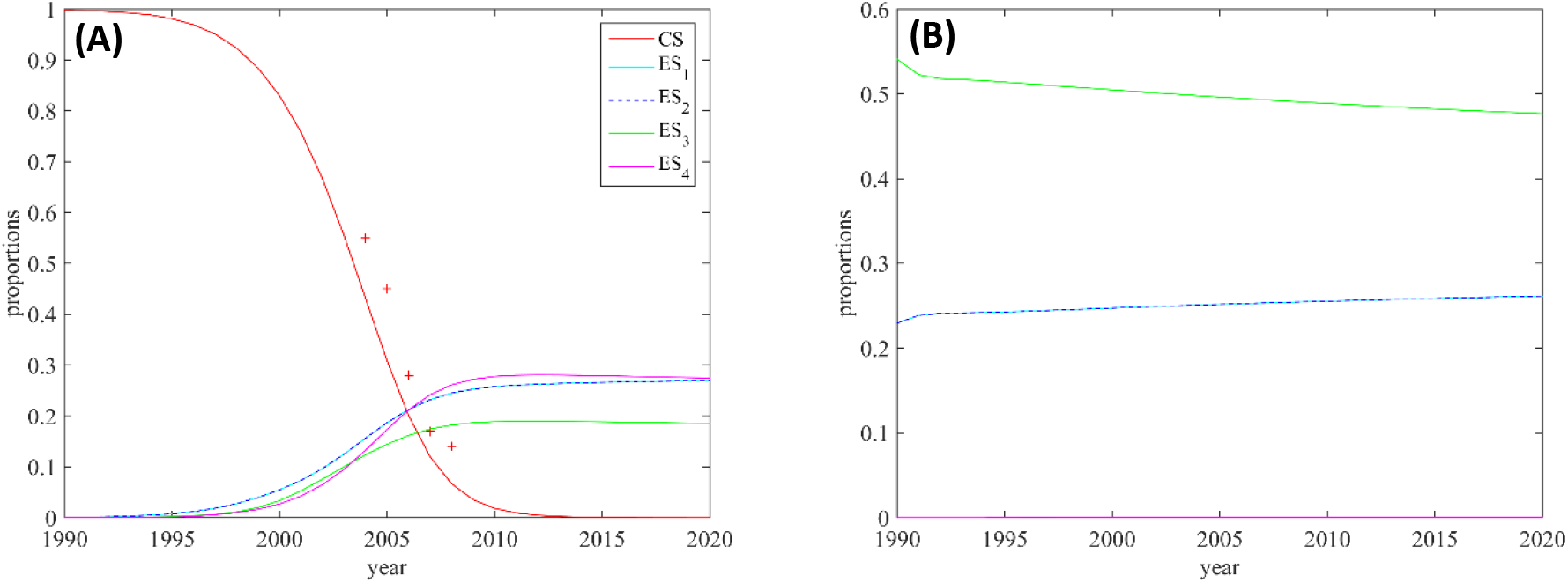
Estimated average proportions of the classical and emerging strains in the study area. *Panel (A):* the plain lines correspond to the simulated proportions and the red crosses correspond to the proportions of CS in the data. *Panel (B):* simulated proportions of the ESs obtained by assuming that the CS is absent. In both cases, the parameter values correspond to the MLE Θ^*^. Note that the curves corresponding to the ESs 1 and 2 are superimposed.

## DISCUSSION

In this work, we developed a reaction-diffusion model to describe the spatial dynamics of invasion of a resident population inhabiting a spatially structured environment by newly introduced variants. Using a mechanistic-statistical framework, we confronted the model to a dataset recording the invasion by several emerging genetic variants of a resident population of WMV and succeeded in (i) estimating the dates and places of successful introduction of each emerging variant as well as parameters related to growth and dispersal, (ii) reconstructing the invasion by the new variants from their introduction sites, (iii) establishing a competitive advantage of the new variants as compared to the resident population and (iv) predicting the fate of each variant. Simulations with the optimal parameter values showed an adequate fit, proving that the model is able to reproduce the observed spatial dynamics despite the strong mechanistic constraints of the model structure and the strong spatial censorship of the dataset, i.e. missing data. We used fraction frequencies in the dataset rather than counting data. We argue that such observations are more robust to heterogeneities in the sampling conditions as they do not require a standardized observation protocol. As stated by [37], abundance data can only be used if the count-proportion, i.e. the ratio between expected count and population size, can safely be assumed to be constant, or if factors affecting variation in the count-proportion can be identified and then accommodated through parametric modeling.

The estimations suggest that three of the four emerging strains have been introduced at approximately the same date, while the fourth one was introduced 5 years later (the model considered only 5-yr intervals of introductions because of constraints on computation time). Despite the neutrality assumption that we made between the emerging strains, we observed different durations of the establishment stage: while ESs 1 and 2 became locally the most prevalent strains only 6 years after their introduction, ES_3_ displayed delayed dynamics since it became locally the most prevalent strain 11 years after its introduction. In comparison, ES_4_ was introduced 5 years later than ES_3_ but became locally the most prevalent strain at about the same date. The low prevalence of the ES_3_ in the dataset could be explained by a lower fitness of this strain, e.g. higher winter decay rate, lower growth rate or weaker competitivity. Our results indicate that this pattern can also be observed with a neutrality assumption, as a result of the joint effects of the local composition of the landscape, and of the position of the sampling sites, far away from the introduction site. Indeed, in the area where ES_4_ was first observed, cucurbits crops are very frequent with high connectivity between crops, whereas the area where ES_3_ was found is patchier. Similarly, our results indicate that the overall prevalence of the CS strain in the study area has been slightly overestimated in the data, due to sampling sites concentrated in the regions where it is indeed the most prevalent strain. The reconstructed dynamics of the five strains therefore underline the importance of estimating jointly the places and dates of the introduction and the other ecological parameters as well as the importance of considering the spatial structure of the sampling design.

Based on WMV-infected samples collected in Southeastern France for more than 30 years, ES_1_ was first detected in 1999 [35] whereas the other ESs were observed in 2002 to 2004 [33], i.e. 9 to 14 years after the estimated introduction dates. Such a lapse between the introduction of a plant pathogen and its first detection is consistent with estimations obtained for other plant viruses [38, 39, 40, 19]. ESs strains have been detected in several European and Mediterranean countries ([14]), and in the USA [41], in the few years following their description in France, and their prevalence in these countries seems to increase even if few time series data are available. The reasons for these almost simultaneous emergences in distant countries and variable environments are not fully understood. WMV being considered so far as not seed-transmitted, the ESs strains have probably been disseminated through long-distance exchanges of plant material [33].

In addition to the dates and places of successful introduction, our model provides estimation of ecological parameters *in natura*. In particular, the diffusion parameter, measuring the mobility of the viruses (or, more precisely, of their aphid vectors) was estimated, leading to a value 0.44 km^2^ per day for WMV. Among plant viruses, there are few estimates of diffusion coefficients, and most estimate rather focus on the speed of range expansion, which can be more directly derived from observations. For instance, an average speed of 33 km/yr and 13 km/yr was estimated for the leafhopper-transmitted wheat streak virus [42] and the whitefly-transmitted East African cassava mosaic virus [43] respectively, to be compared with the spread rate of 5 km/year found here. For non-persistently aphid-transmitted viruses like WMV and the other potyviruses, the insect remains viruliferous after probing on an infected plant for only a few minutes to hours [14], and the dispersal distance is supposed to be limited, even if in exceptional climatic conditions, the potyvirus maize dwarf mosaic virus was transmitted by viruliferous aphids over more than 1000 km [44]. Estimations for the potyvirus plum pox virus indicated that 50% of the infectious aphids leaving an infected plant land within about 90 meters, while about 10% of flights terminate beyond 1 km [45]. Here, a diffusion coefficient of 0.44 km^2^ per day corresponds to a mean dispersal distance of 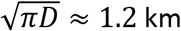 after 1 day (see Supplementary Note S7), which seems in agreement with these data.

Another critical parameter that was estimated is the winter decay rate. Most studies focus on the epidemic period but off-season dynamics can be crucial to understand demography and genetic diversity [32, 46]. We assumed here that emerging strains differed from the resident population only through this parameter. Indeed, no obvious differences in host range, including both cultivated hosts and weeds, have been found between CS and ES strains, whereas ESs strains were found to be better transmitted than CS ones from some weeds infected by both CS and ESs [47]. This could contribute to more efficient transfer from weeds to crops at the beginning of the growing season, leading to a lower winter decay of ESs strains. In our modelling approach we found that ESs strains were able to invade the resident population because of a lower decay during winter. Nevertheless, as all of the other parameters in the model of population dynamics have been set to the same value for the CS and the ESs, all of the competitive advantage of the ESs can be expressed only through the decay rates, explaining these differences, and the unrealistic 0-decay rate of the ESs. The unrealistic null winter decay rate that we found for the ESs suggests that, contrarily to our assumptions, competitive advantage of ESs probably occurs also during summer. Consistently, there is no efficient cross-protection between CS and ESs strains [47], but [21] found that superinfection by ESs strains of a plant already infected by CS is easier than the opposite situation. In our model, we also make a neutrality assumption between the ESs that differ only by their introduction dates/sites. This assumption may not completely reflect the complexity of the interactions between viral strains and the biological variability between and within molecular groups [47].

In 2008, the ESs have replaced the CS, which was not anymore the most prevalent strain, whatever the position in the study area, 18 years after the first ESs introduction. Moreover, this competitive advantage of the ESs is expected to lead to the total replacement of the CS by the ESs in about 25 years, i.e. in 2015. These results are consistent with current knowledge: new observations carried out in 2016 and 2017 showed than the classical CS strain is no more detectable [48]. Besides the disappearance of CS strains, the surveys performed in 2016-2017 revealed a complex and dynamic situation that fitted partially with the model. ES_3_ was detected in only one location in the southwestern part of the study area, confirming its low dispersal and probable low fitness. As predicted by the model, ES_1_ and ES_2_ were present in the Eastern part of the area and ES_4_ was present in all the study area. However, it was found to be more prevalent than ES_1_ and ES_2_ even in the Eastern part, suggesting that its fitness is higher than ES_1_ and ES_2_. New variants, not detected in 2004-2008, were also observed in 2016-2017, and some of them presented a high prevalence in all the area. Deep sequencing of two genomic regions revealed, contrary to the 2004-2008 situation [49], a high prevalence of recombinants among these new strains [48], blurring the distinction between molecular groups based on CP sequences only. As in 2004-2008, landscape heterogeneity seemed to affect virus dispersal [48].

In a more general perspective, this study shows how mechanistic approaches can be used to infer the historical dynamics of invasive genotypes or species from initial introduction. These approaches enable considering hypothetical scenarios, to get a better understanding of the impact of the biological interactions on the overall dynamics. Here, in agreement with theoretical results in [50], the simulations without the CS strain showed that its presence promotes a higher diversity among the emerging strains, by altering the outcome of the competition between the ESs, and by slowing down the overall dynamics, thus reducing founder effects. Another advantage of using a mechanistic-statistical approach, compared to a correlative approach, is that the parameter values bring some insight into biological processes and life history traits, (e.g. the diffusion coefficient is related to dispersal ability). Good knowledge of the parameter values, especially for the biological parameters, will be helpful for future modelling, either with reaction-diffusion models or with other approaches such as stochastic diffusion models, which share some common parameters with reaction-diffusion models (e.g., [10]. The method developed in this work is computationally very costly. We plan to develop much faster methods, based on deterministic optimization algorithms and analytic descriptions of the gradients of the likelihood. Introducing specific fitness parameters, besides winter decay rate, for the different groups will also help to better understand the effect of interactions between variants on the evolution of viral populations.

## ACKNOWLEDGEMENTS

This work was also funded by INRAE grant “MEDIA”.

## AUTHOR CONTRIBUTIONS STATEMENT

All authors conceived the model; L.R. and J.P. wrote the first draft; L.R. carried out the numerical computations; all authors discussed the results and contributed to the final manuscript.

## SUPPLEMENTARY INFORMATION

### Supplementary Fig. S1. Effect of the initial population density

We computed the proportions of each strain with the MLE Θ^*^, assuming other initial densities of the ESs instead of 1/10^th^ of the carrying capacity at the introduction point. **Fig. S1** depicts the proportions of each strain, assuming either that the initial density of the ESs are increased or decreased by a factor 2.

**Fig. S1.**
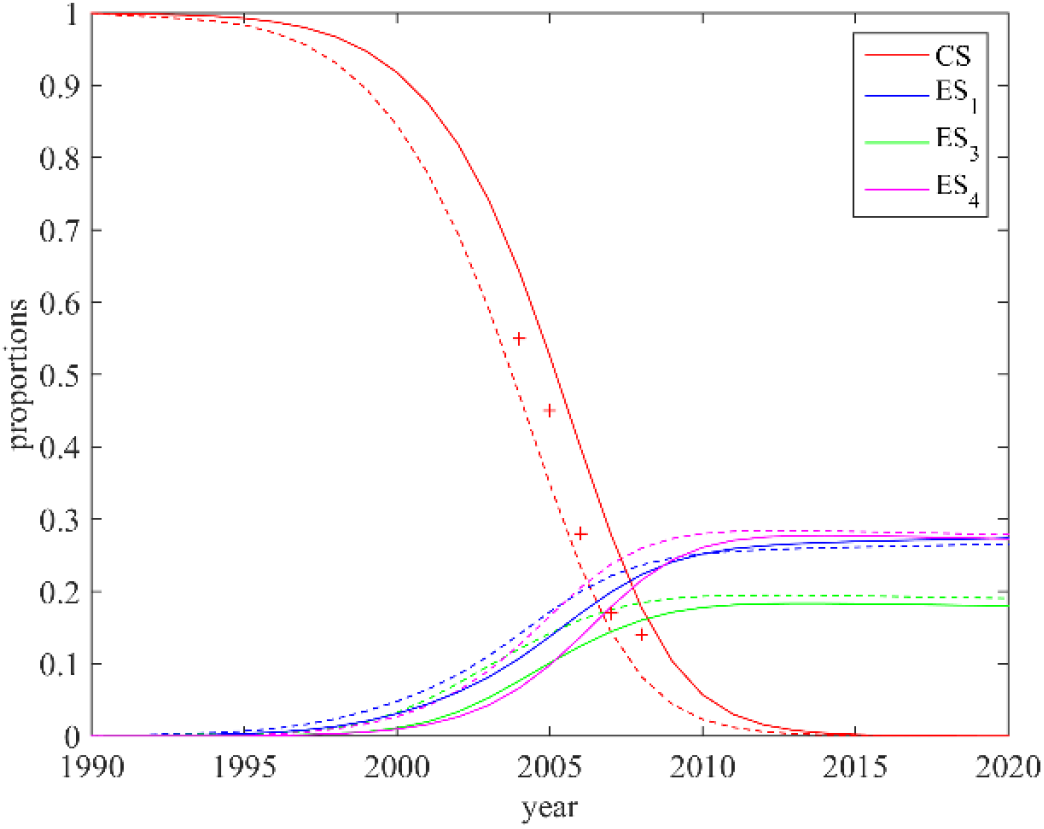
Estimated average proportions of the classical and emerging strains in the study area with varying values of the initial density of the ESs. The plain lines correspond to initial densities divided by 2 (1/20^th^ of the carrying capacity) and the dotted lines to initial densities multiplied by 2 (1/5^th^ of the carrying capacity).

### Supplementary Fig. S2. Finite element method

Partial differential equations were solved with Comsol Multiphysics^®^ time-dependent solver which is based on a finite element method (FEM). The triangular mesh which was used for our computations is depicted below (**Fig. S2**). It is made of 4706 triangular elements.

**Fig. S2.**
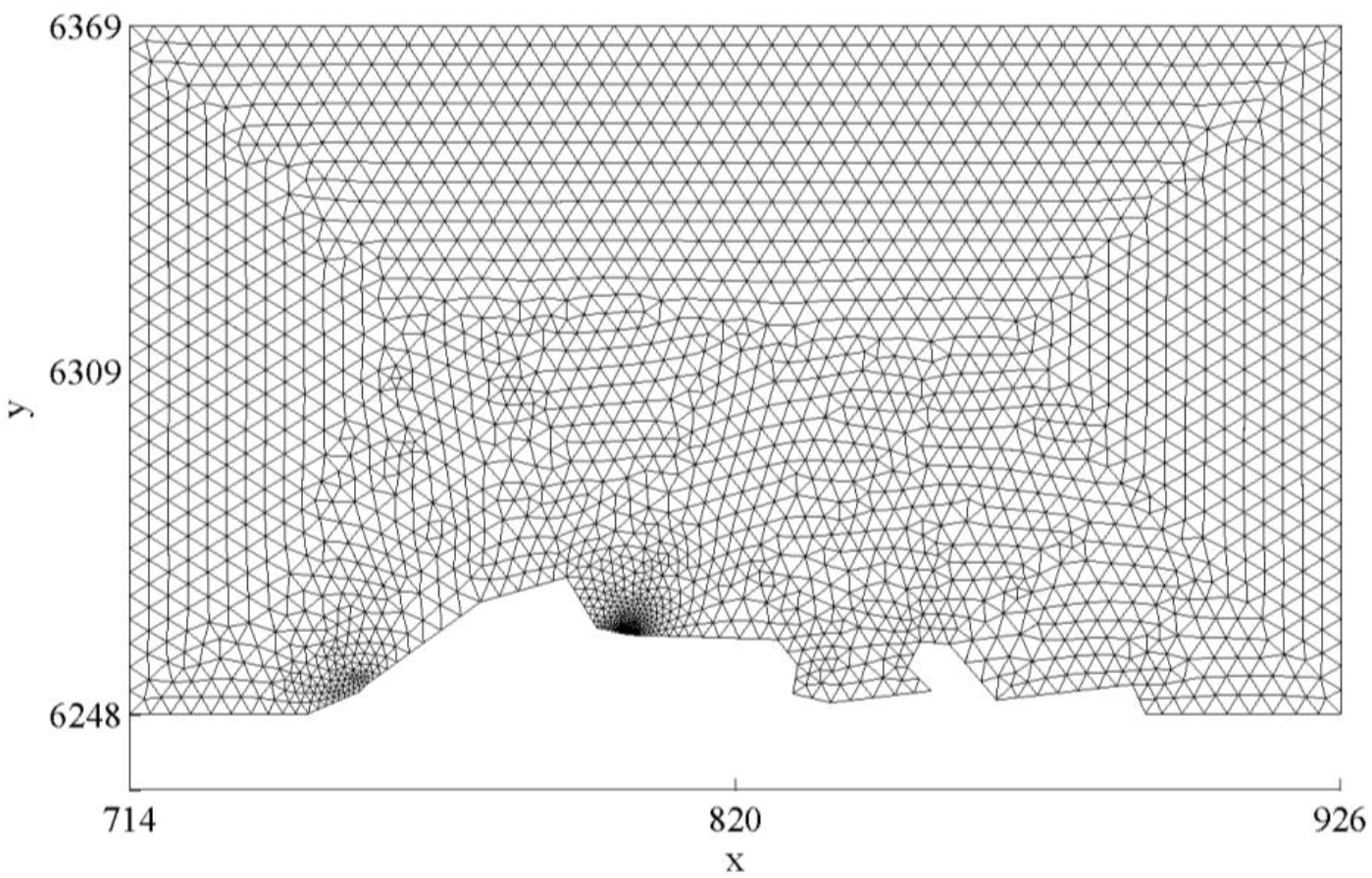
Triangular mesh of the study site.

### Supplementary Fig. S3: Introduction points

**Fig. S3.**
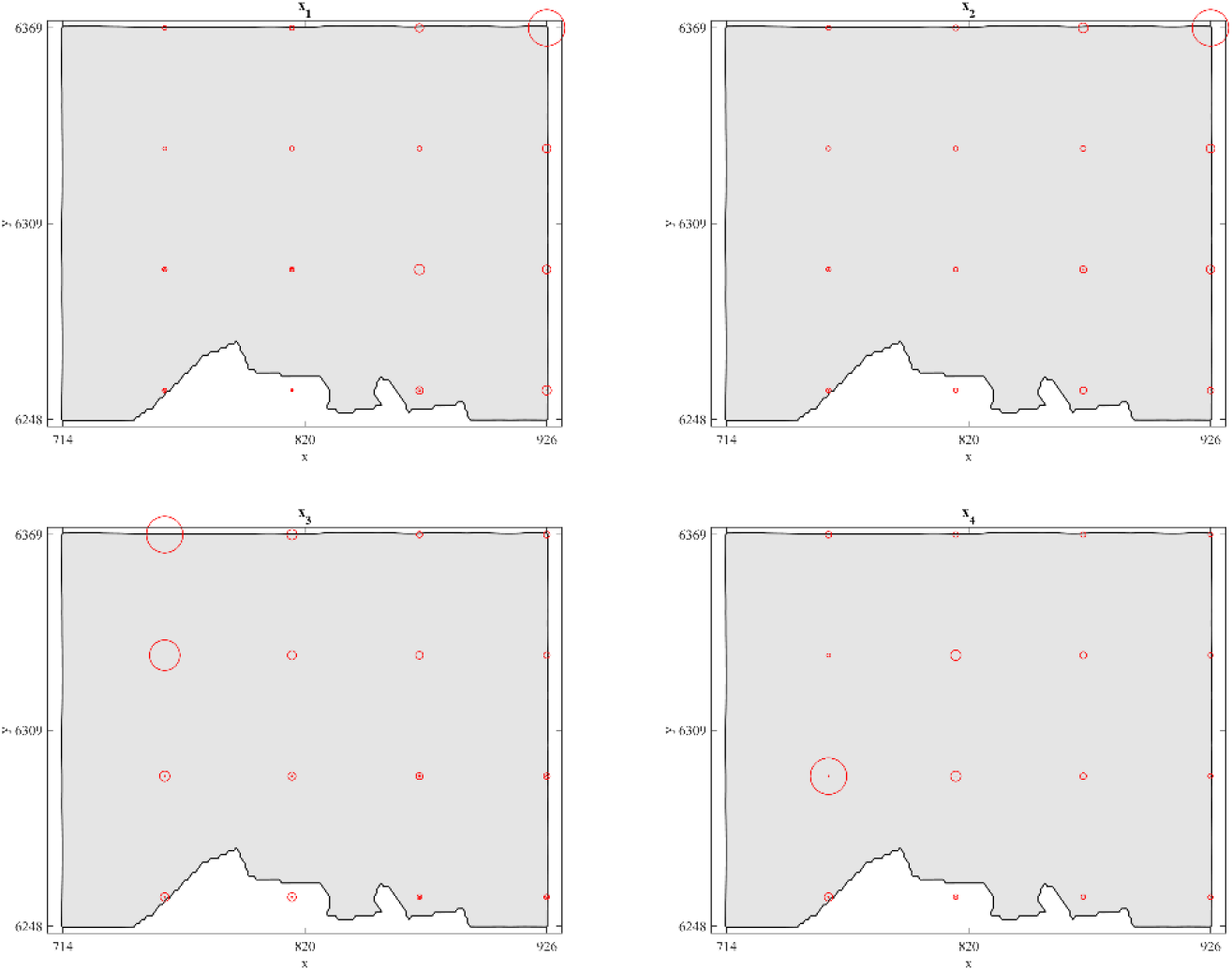
Likelihood function in terms of the introduction point. The area of the circles are proportional to the highest value reached by the function *f*(Θ_*j*_) when the introduction point of the *ES*_*k*_ (each panel corresponds to a different strain) is located at the center of the circle.

### Supplementary Fig. S4: Profile of the likelihood function

The simulated annealing algorithms led to 6 sequences 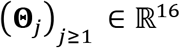, for a total of 32000 elements Θ_*j*_ and 32000 evaluations of 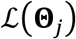. Fig. S4 depicts the values of a monotone transform of the likelihood 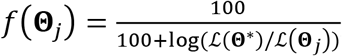, projected onto the 1D variables *D*, *r*, *m_c_*, *m*_*e*_, *n*_1_, *n*_2_, *n*_3_, *n*_4_. We observe that close parameter values tend to lead to close values of the likelihood, which strongly suggests that the MLE Θ* is close to the actual maximizer of the likelihood function.

**Fig. S4.**
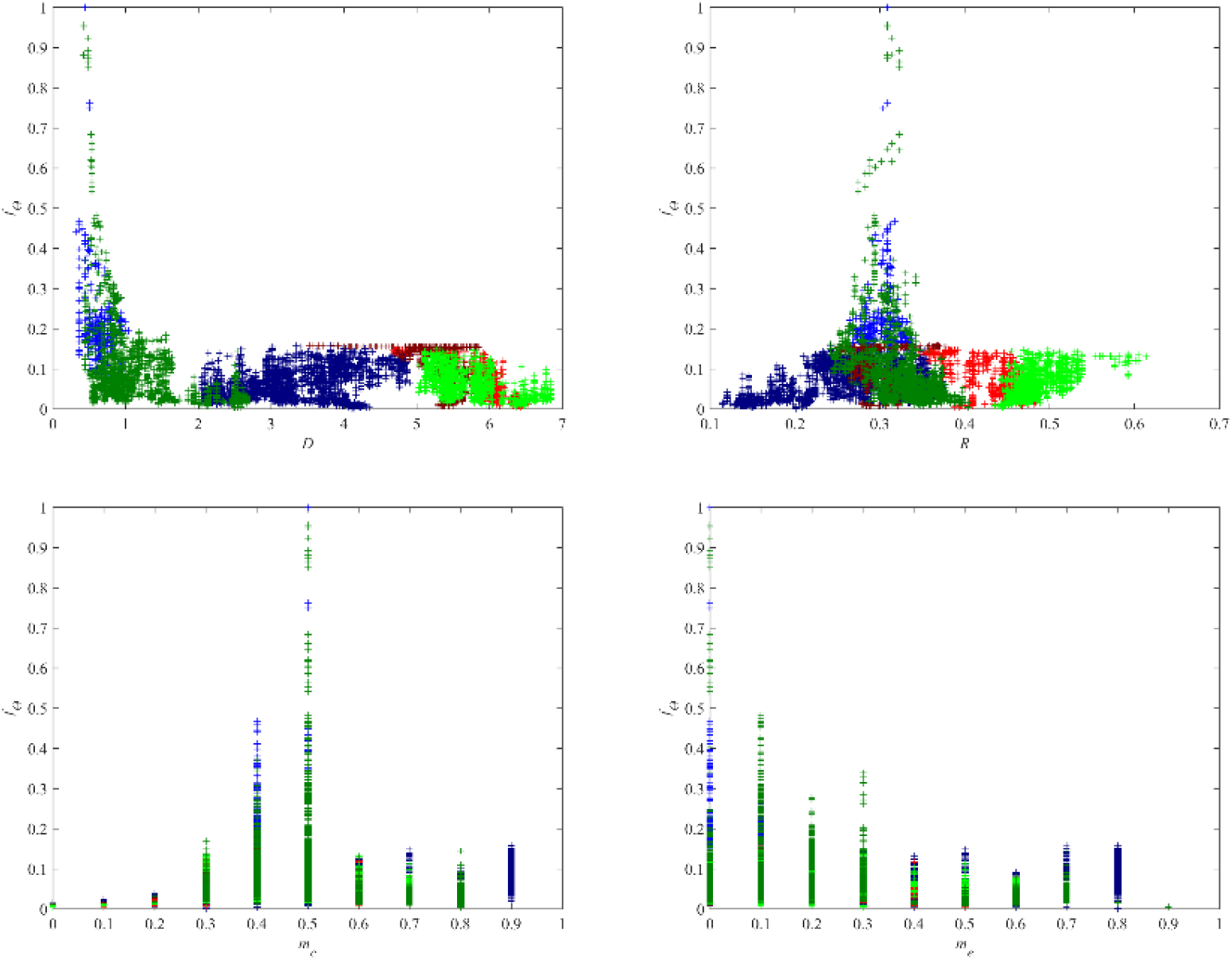
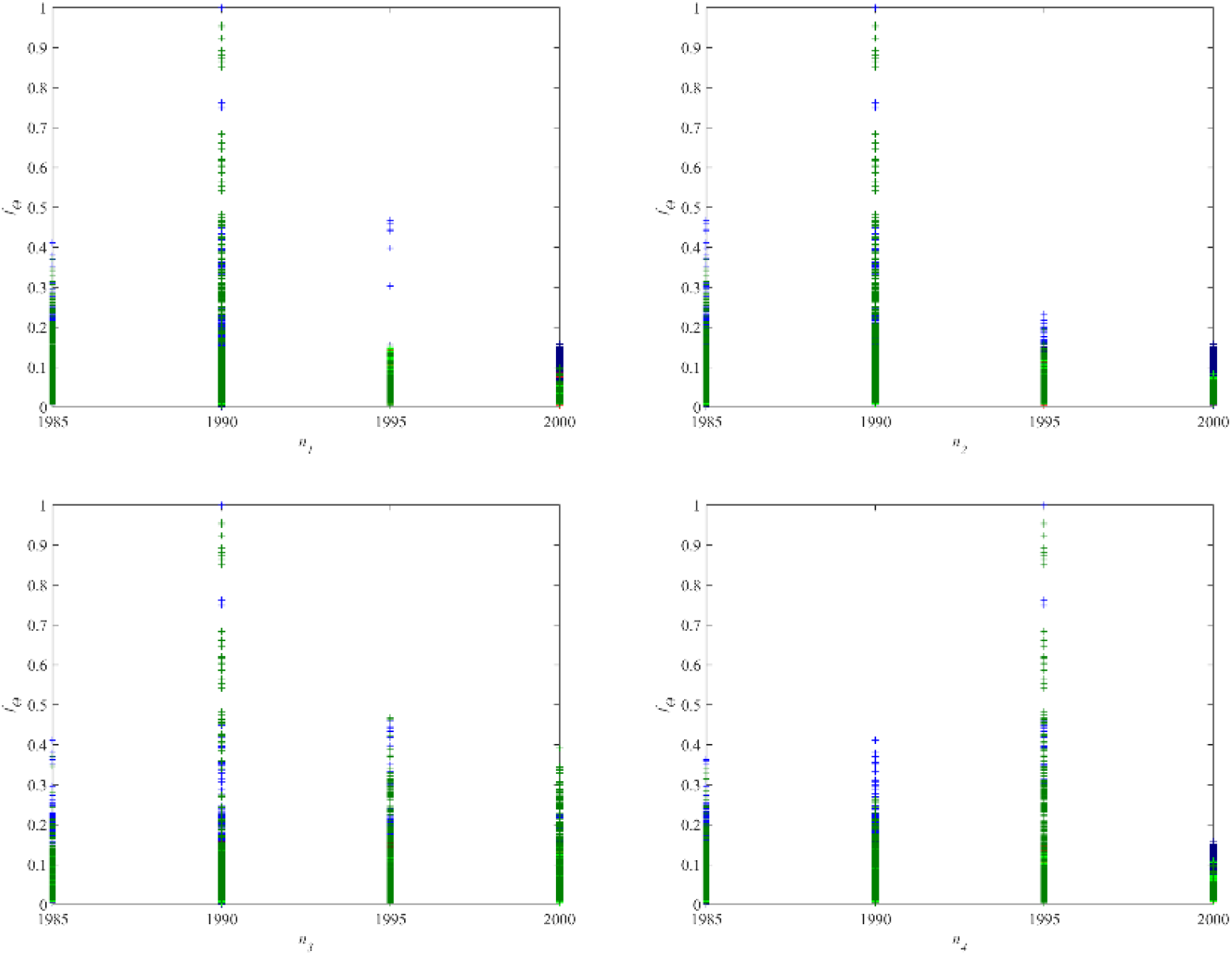
Profile of the likelihood function. Each panel corresponds to a projection of the 32000 computed values of *f*(Θ_*j*_), over the 1D variables *D*, *r*, *m*_*c*_, *m*_*e*_, *n*_1_, *n*_2_, *n*_3_, *n*_4_, respectively. The blue crosses corresponds to parameters Θ_*j*_ obtained with the slowest cooling rate (*α* = 0.9995, 2 chains); the green crosses to the intermediate cooling rate (*α* = 0.999, 2 chains) and the red crosses correspond to fastest cooling rate (*α* = 0.995, 2 chains). Note that *f*(Θ^*^) = 1 and *f*(Θ) = 0.1 when 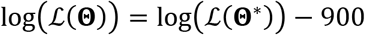 (here, 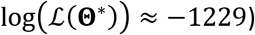).

### Supplementary Fig. S5: Proportions of the classical and emerging strains in the landscape: 2008

**Fig. S5.**
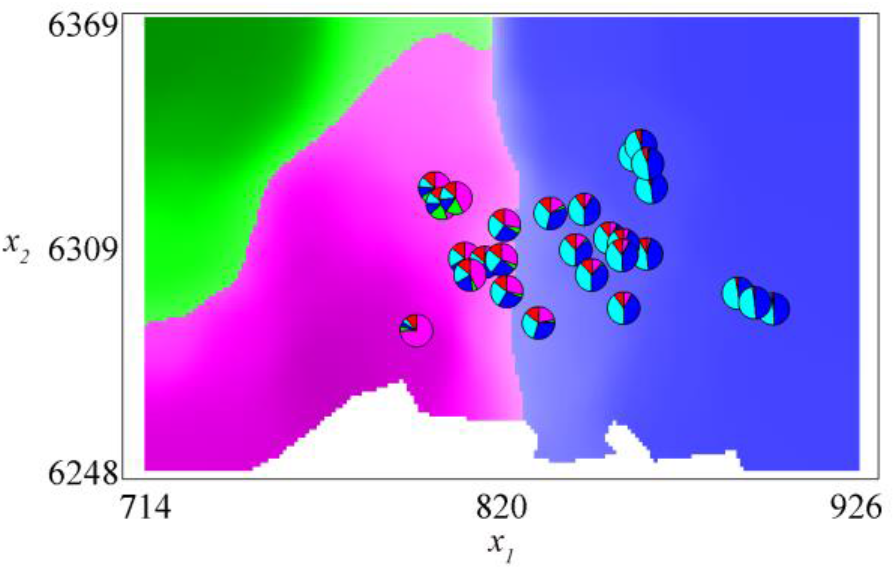
Proportions of the classical and emerging strains in the landscape: data and simulations. The colors of the shaded regions indicate which strain is the most prevalent. The red regions correspond to the classical strain; light blue and blue: *ES*_1_, *ES*_2_ (these two strains have the same density, only *ES*_2_ is represented); green: *ES*_3_; pink: *ES*_4_. The pie charts describe the relative proportions of the strains found in the data (same color legend). The simulation results presented here correspond to the middle of the intra-annual stage (2^nd^ week of June), and were obtained with the MLE Θ^*^.

### Supplementary Fig. S6. Full timeline of the dynamics of the different strain proportions in the landscape, obtained with the maximum likelihood estimate (MLE)Θ^*^

The pictures below represent the dynamics of the classical and emerging strains, from the first estimated introduction date of an emerging strain (1990) to 2019. The colors of the shaded regions indicate which strain is the most prevalent. The red regions correspond to the classical strain; blue: ES_1_, ES_2_ (these two strains have the same density, only ES_1_ is represented); green: ES_3_; pink: ES_4_. The results presented here correspond to the middle of the intra-annual stage (2^nd^ week of June).

**Fig. S6.**
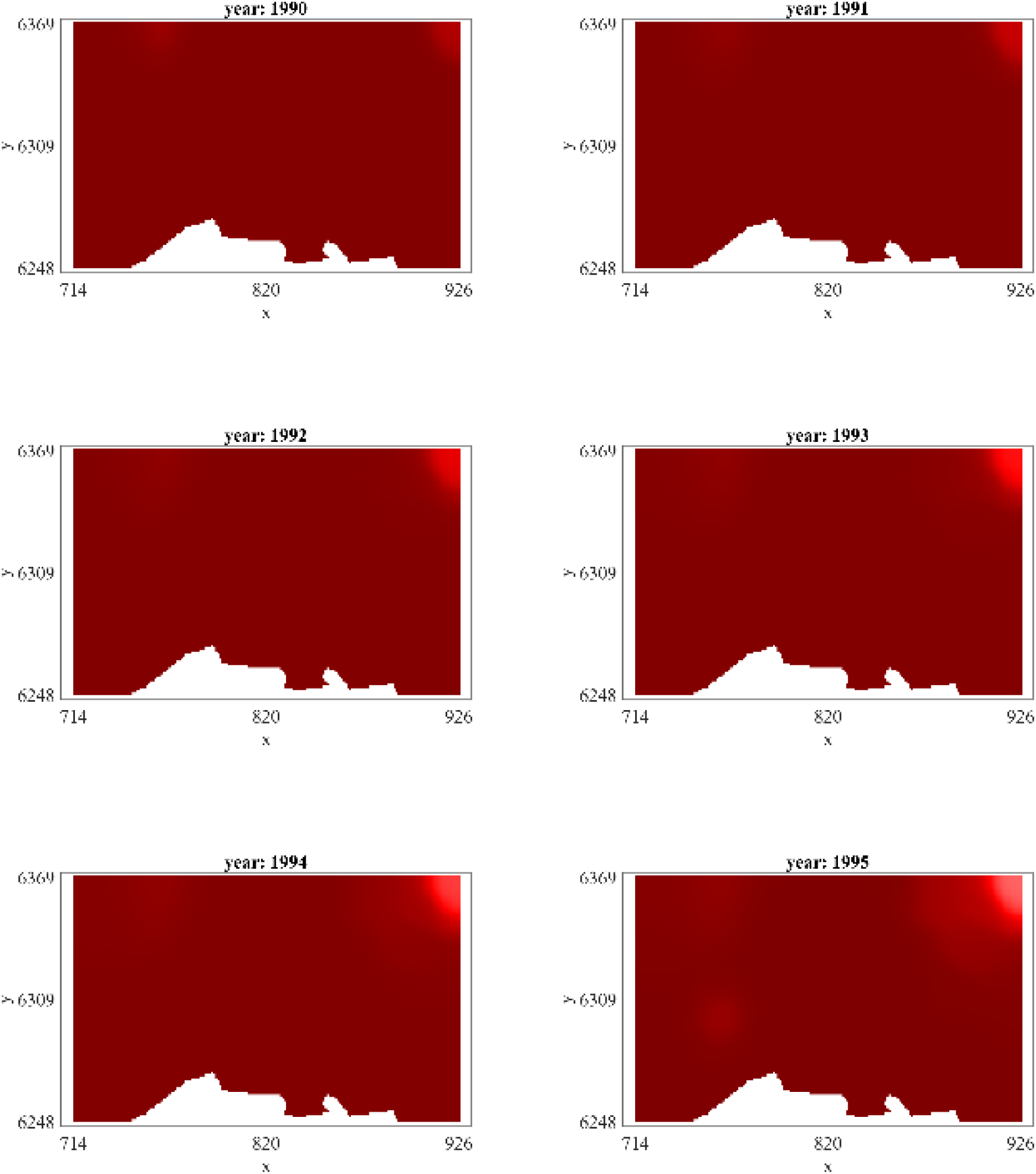
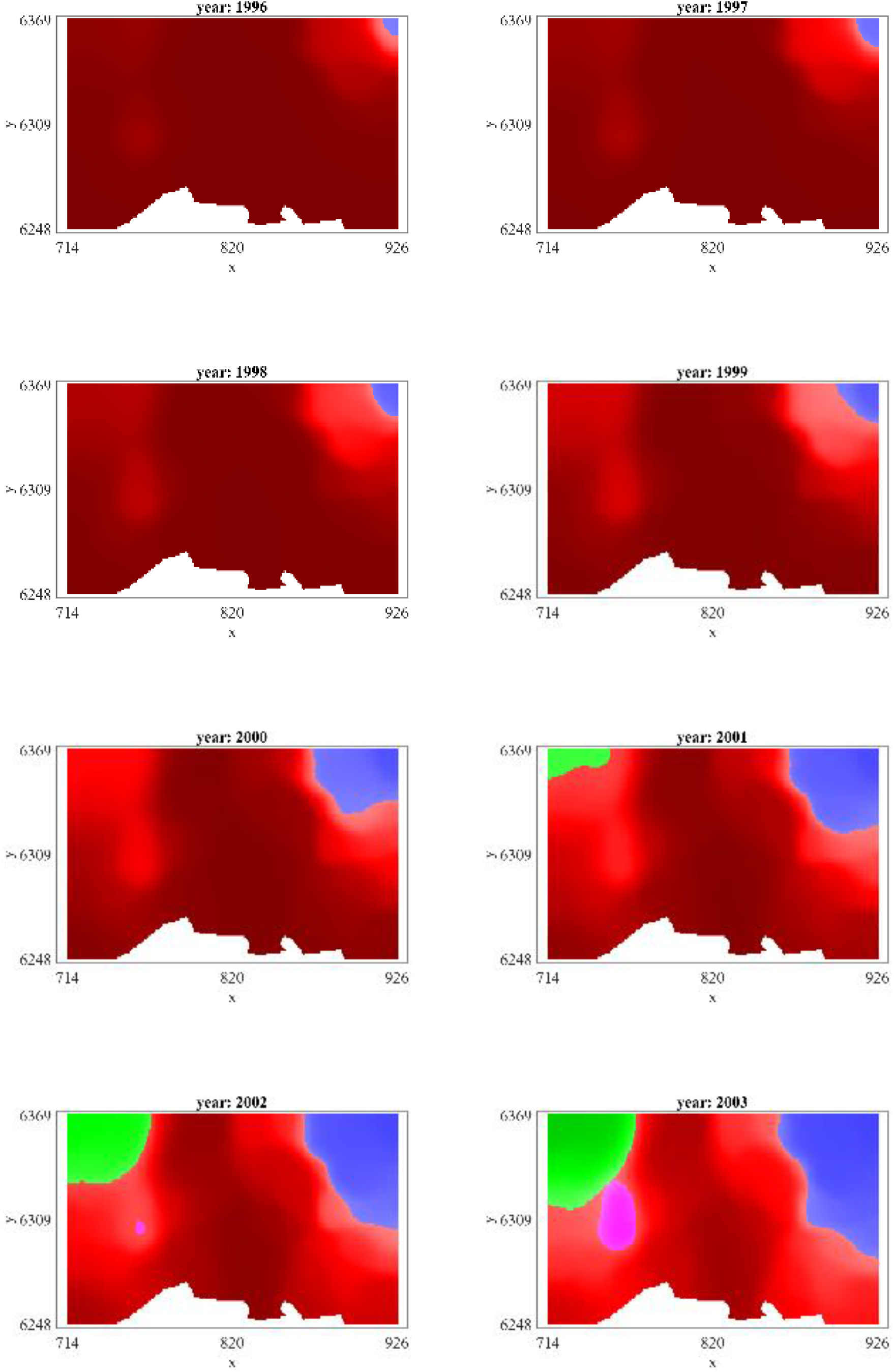
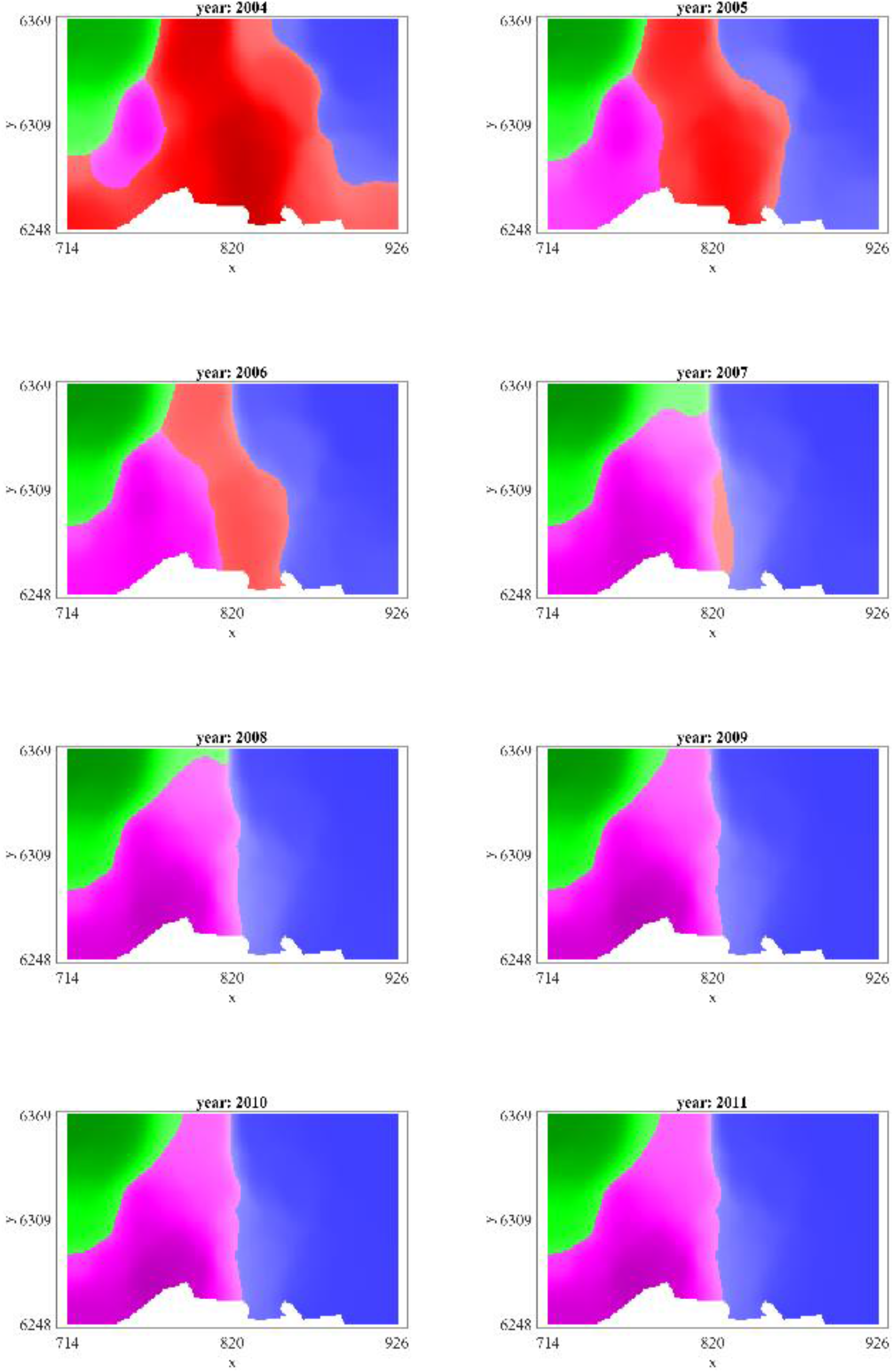
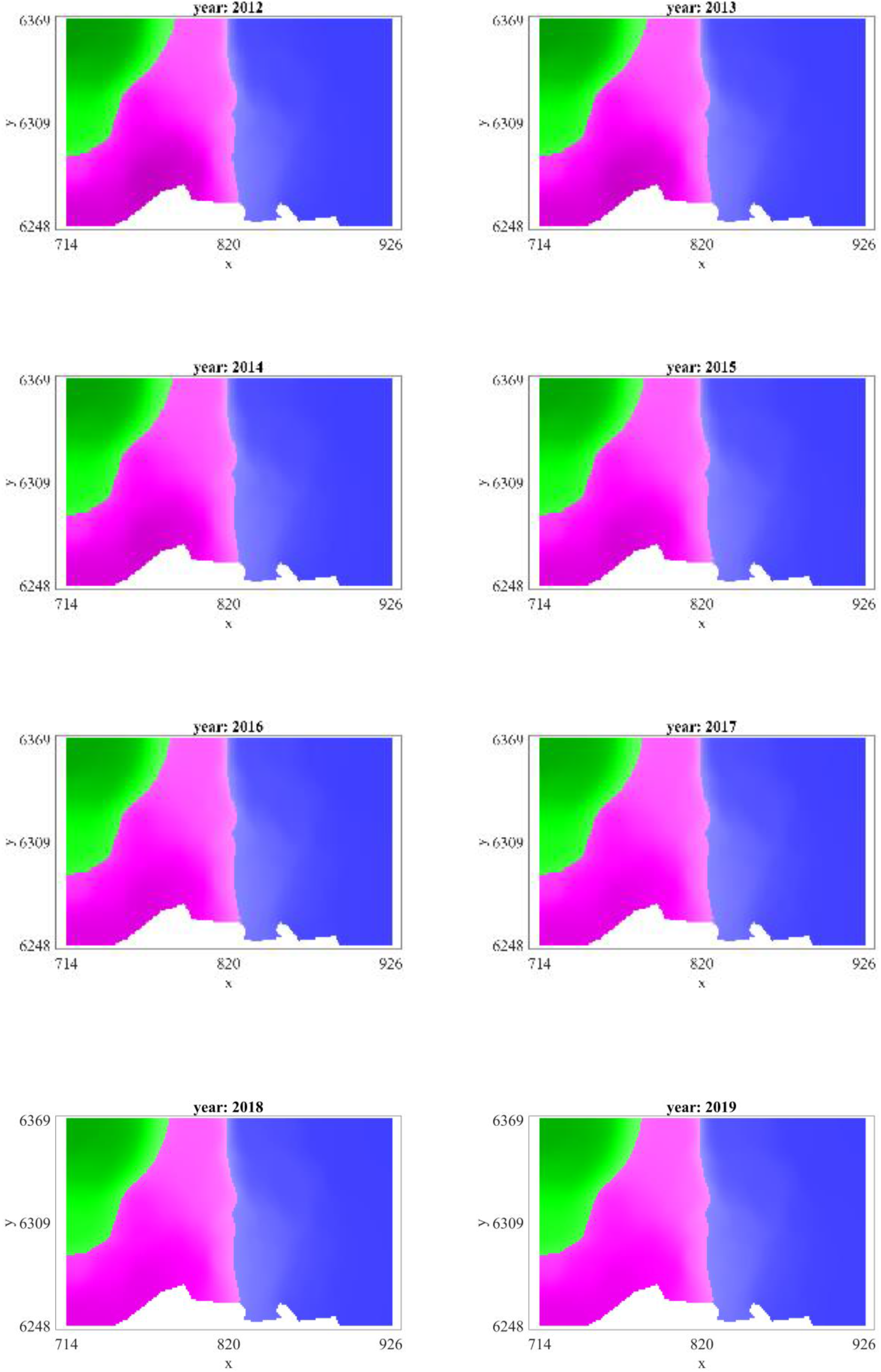

### Supplementary Note S7. A priori bounds on the parameter values, and relationship between diffusion coefficient and dispersal distance

Assuming a random walk movement with discrete space step *λ* and time step *т*, the corresponding diffusion coefficient is 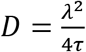. With a time step *т* equal to 1 day, we get 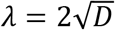. The bounds *D* ∈ (10^−4^, 10) km^2^/day thus correspond to space steps 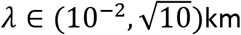. In other words, each day the average distance travelled by a virus (through its vector) is *a priori* assumed to be comprised between 10 m and 3.16 km.

The solution of a pure diffusion equation (i.e., without reproduction), *∂*_*t*_*u*(*t*, ***x***) = *D*Δ*u* in dimension 2, starting from a localized initial condition at ***x*** = 0 is 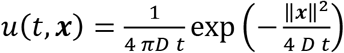. The mean dispersal distance after 1 day is:

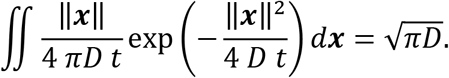

Having a growth rate *r* day^−1^ means an increase by a factor *e*^*r*^ each day, in the absence of competition. The bounds *r* = 0.1 and *r* = 1 thus correspond a daily increase by a factor comprised between 1.1 and 2.7.

### Supplementary Note S8. Computation of likelihood-ratio based confidence intervals

To compute confidence intervals for *θ*_*i*_, where *Θ* = (*Θ*_1_, …, *Θ*_16_), we first define the profile likelihood function, 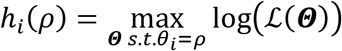. The (1 − α) confidence intervals for *Θ*_*i*_ can be constructed by finding the set of parameter values *p* such that 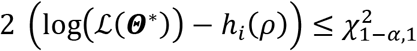, where 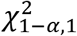 is the (1 − α) percentile of the χ^2^ distribution with 1 degree of freedom [1] (note that 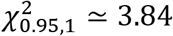). For each value of *p*, *h*_*i*_(*p*) was computed based on the values of *Θ*_*j*_ which have been explored by the simulated annealing algorithm (using the 6 sequences together, this corresponds to ≈ 32000 values for *Θ*_*j*_). For the parameters *D*, *r* which have not been discretized, to be able to compute *h*_*i*_(*p*) for values of *p* which have not been explored by the algorithm, we approached 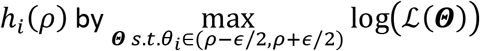, for some *∈* ≥ 0 (we took 1/100^th^ of the length of the support of the prior distribution). For the other parameters, *h*_*i*_(*p*) is computed by interpolation between the discrete grid points.

## REFERENCES

[1] Hanno Seebens, Tim M Blackburn, Ellie E Dyer, Piero Genovesi, Philip E Hulme, Jonathan M Jeschke, Shyama Pagad, Petr Pyšek, Marten Winter, Margarita Arianoutsou, et al. No saturation in the accumulation of alien species worldwide. Nature communications, 8(1):1–9, 2017.

[2] Raghavan Charudattan. Biological control of weeds by means of plant pathogens: significance for integrated weed management in modern agro-ecology. BioControl, 46(2):229–260, 2001.

[3] Todd A Crowl, Thomas O Crist, Robert R Parmenter, Gary Belovsky, and Ariel E Lugo. The spread of invasive species and infectious disease as drivers of ecosystem change. Frontiers in Ecology and the Environment, 6(5):238–246, 2008.

[4] A K Sakai, F W Allendorf, J S Holt, D M Lodge, J Molofsky, K A With, S Baughman, R J Cabin, J E Cohen, N C Ellstrand, D E Mccauley, P O’Neil, I M Parker, J N Thompson, and S G Weller. The population biology of invasive species. Annual Review Of Ecology And Systematics, 32:305–332, 2001.

[5] D Martinetti and S Soubeyrand. Identifying lookouts for epidemio-surveillance: Application to the emergence of Xylella fastidiosa in france. Phytopathology, 109(2):265–276, Feb 2019.

[6] S F Elena, A Fraile, and F Garcia-Arenal. Evolution and emergence of plant viruses. In K Maramorosch and Fa Murphy, editors, Advances In Virus Research, Vol 88, volume 88 of Advances In Virus Research, pages 161–191. 2014.

[7] M J Mcleish, A Fraile, and F Garcia-Arenal. Trends and gaps in forecasting plant virus disease risk. Annals Of Applied Biology, 176(2):102–108, Mar 2020.

[8] L Rimbaud, C Bruchou, S Dallot, D R J Pleydell, E Jacquot, S Soubeyrand, and G Thebaud. Using sensitivity analysis to identify key factors for the propagation of a plant epidemic. Royal Society Open Science, 5(1), Jan 2018.

[9] C Abboud, O Bonnefon, E Parent, and S Soubeyrand. Dating and localizing an invasion from post-introduction data and a coupled reaction–diffusion–absorption model. Journal of Mathematical Biology, 79(2):765–789, 2019.

[10] L Roques, E Walker, P Franck, S Soubeyrand, and E K Klein. Using genetic data to estimate diffusion rates in heterogeneous landscapes. Journal of mathematical biology, 73(2):397–422, 2016.

[11] S Soubeyrand, P De Jerphanion, O Martin, M Saussac, C Manceau, P Hendrikx, and C Lannou. Inferring pathogen dynamics from temporal count data: The emergence of Xylella fastidiosa in france is probably not recent. New Phytologist, 219(2):824–836, Jul 2018

[12] P Lefeuvre, D P Martin, G Harkins, P Lemey, A J A Gray, S Meredith, F Lakay, A Monjane, J-M Lett, A Varsani, and J Heydarnejad. The spread of tomato yellow leaf curl virus from the middle east to the world. Plos Pathogens, 6(10), Oct 2010.

[13] E Moriones, S Praveen, and S Chakraborty. Tomato leaf curl New Delhi virus: An emerging virus complex threatening vegetable and fiber crops. Viruses-Basel, 9(10), Oct 2017.

[14] H Lecoq and C Desbiez. Viruses of cucurbit crops in the mediterranean region: An ever-changing picture. In G Loebenstein and H Lecoq, editors, Viruses And Virus Diseases Of Vegetables In The Mediterranean Basin, volume 84 of Advances In Virus Research, pages 67–126. 2012.

[15] C Desbiez, B Moury, and H Lecoq. The hallmarks of “green” viruses: Do plant viruses evolve differently from the others? Infection Genetics And Evolution, 11(5):812–824, Jul 2011

[16] A J Drummond, O G Pybus, A Rambaut, R Forsberg, and A G Rodrigo. Measurably evolving populations. Trends In Ecology & Evolution, 18(9):481–488, Sep 2003.

[17] P Lemey, A Rambaut, A J Drummond, and M A Suchard. Bayesian phylogeography and finds its. Plos Computational Biology, 5(9), Sep 2009.

[18] O G Pybus, M A Suchard, P Lemey, F J Bernardin, A Rambaut, F W Crawford, R R Gray, N Arinaminpathy, S L Stramer, M P Busch, and E L Delwart. Unifying the spatial epidemiology and molecular evolution and of emerging epidemics. Proceedings Of The National Academy Of Sciences Of The United States Of America, 109(37):15066–15071, Sep 11 2012.

[19] M Rakotomalala, B Vrancken, A Pinel-Galzi, P Ramavovololona, E Hebrard, J S Randrianangaly, S Dellicour, P Lemey, and D Fargette. Comparing patterns and scales of plant virus phylogeography: Rice yellow mottle virus in madagascar and in continental africa. Virus Evolution, 5(2), Jul 2019.

[20] S Soubeyrand and L Roques. Parameter estimation for reaction-diffusion models and of biological invasions. Population Ecology, 56(2):427–434, 2014.

[21] F Fabre, J Chadoeuf, C Costa, H Lecoq, and C Desbiez. Asymmetrical over-infection as a process of and plant virus. Journal Of Theoretical Biology, 265(3):377–388, Aug 7 2010.

[22] F Perefarres, G Thebaud, P Lefeuvre, F Chiroleu, L Rimbaud, M Hoareau, B Reynaud, and J-M Lett. Frequency-dependent assistance as a way out of competitive exclusion between two strains of an emerging virus. Proceedings Of The Royal Society B-Biological Sciences, 281(1781), Apr 22 2014.

[23] Ana Beatriz M and J Lopez-Moya. When viruses play team sports: Mixed and infections in plants. Phytopathology, 110(1):29–48, Jan 2020

[24] P Turchin. Quantitative Analysis of Movement: Measuring and Modeling Population Redistribution in Animals and Plants. Sinauer, Sunderland, MA, 1998.

[25] Alan Hastings, Kim Cuddington, Kendi F Davies, Christopher J Dugaw, Sarah Elmendorf, Amy Freestone, Susan Harrison, Matthew Holland, John Lambrinos, Urmila Malvadkar, and et al. The spatial spread of invasions: new developments in and theory and evidence. Ecol Lett, 8(1):91–101, 2005.

[26] N Shigesada and K Kawasaki. Biological Invasions: Theory and Practice. Oxford Series in Ecology and Evolution, Oxford: Oxford University Press, 1997.

[27] H Berestycki, F Hamel, and L Roques. Analysis of the periodically fragmented environment model: I - Species persistence. J. Math. Biol., 51(1):75–113, 2005.

[28] L Roques, S Soubeyrand, and J Rousselet. A statistical-reaction-diffusion approach for and analyzing expansion. Journal Of Theoretical Biology, 274(1):43–51, Apr 7 2011.

[29] L M Berliner. Physical-statistical and modeling in geophysics. J Geophys Res, 108:8776, 2003.

[30] S Soubeyrand, A-L Laine, I Hanski, and A Penttinen. Spatiotemporal structure of host-pathogen interactions and in a metapopulation. American Naturalist, 174(3):308–320, Sep 2009.

[31] S Soubeyrand, S Neuvonen, and A Penttinen. Mechanical-statistical modeling in ecology: from outbreak detections and to pest dynamics. Bull Math Biol, 71:318–338, 2009.

[32] H Lecoq, C Wipf-Scheibel, K Nozeran, P Millot, and A Penttinen. Comparative molecular epidemiology provides new insights into zucchini yellow mosaic virus and occurrence in france. Virus Research, 186(Si):135–143, Jun 24 2014.

[33] C Desbiez, B Joannon, C Wipf-Scheibel, C Chandeysson, and H Lecoq. Emergence of new strains of watermelon mosaic virus in south-eastern france: Evidence for limited spread but rapid and local population shift. Virus Research, 141(2, Si):201–208, May 2009.

[34] J D Murray. Mathematical Biology. Third edition, Interdisciplinary Applied Mathematics 17, Springer-Verlag, New York, 2002.

[35] C Desbiez, C Costa, C Wipf-Scheibel, M Girard, and H Lecoq. Serological and molecular variability of watermelon mosaic virus (genus Potyvirus). Archives Of Virology, 152(4):775–781, 2007.

[36] S Kirkpatrick, C D Gelatt, and M P Vecchi. Optimization and by simulated. Science, 220:671–680, 1983.

[37] William A Link and John R Sauer. Estimating relative abundance and from count. Austrian Journal of Statistics, 27(1&2):83–97, 1998.

[38] Gordon W Harkins, Darren P Martin, Siobain Duffy, Aderito L Monjane, Dionne N Shepherd, Oliver P Windram, Betty E Owor, Lara Donaldson, Tania Van Antwerpen, Rizwan A Sayed, et al. Dating the origins of the maize-adapted strain of maize streak virus, MSV-A. The Journal of general virology, 90(Pt 12):3066, 2009.

[39] Ryosuke Yasaka, Hirofumi Fukagawa, Mutsumi Ikematsu, Hiroko Soda, Savas Korkmaz, Alireza Golnaraghi, Nikolaos Katis, Simon YW Ho, Adrian J Gibbs, and Kazusato Ohshima. The timescale of emergence and spread of and turnip mosaic potyvirus. Scientific reports, 7(1):1–14, 2017.

[40] Mohammad Hajizadeh, Adrian J Gibbs, Fahimeh Amirnia, and Miroslav Glasa. The global phylogeny of plum pox and virus is emerging. Journal of General Virology, 100(10):1457–1468, 2019.

[41] Paul Vincelli and Kenneth Seebold. Report of a watermelon mosaic potyvirus strain in Kentucky and undetected by ELISA. Plant Health Progress, 10(1):47, 2009.

[42] A L Monjane, G W Harkins, D P Martin, P Lemey, P Lefeuvre, D N Shepherd, S Oluwafemi, M Simuyandi, I Zinga, E K Komba, D P Lakoutene, N Mandakombo, J Mboukoulida, S Semballa, A Tagne, F Tiendrebeogo, J B Erdmann, T Van Antwerpen, B E Owor, B Flett, M Ramusi, O P Windram, R Syed, J-M Lett, R W Briddon, P G Markham, E P Rybicki, and A Varsani. Reconstructing the history of maize streak virus strain a dispersal to reveal diversification hot spots and its origin and in southern Africa. Journal Of Virology, 85(18):9623–9636, Sep 2011.

[43] Alexandre De Bruyn, Julie Villemot, Pierre Lefeuvre, Emilie Villar, Murielle Hoareau, Mireille Harimalala, Anli L Abdoul-Karime, Chadhouliati Abdou-Chakour, Bernard Reynaud, Gordon W Harkins, et al. East African cassava mosaic-like viruses from Africa to Indian ocean islands: molecular diversity, evolutionary history and geographical dissemination of and a bipartite begomovirus. BMC evolutionary biology, 12(1):228, 2012.

[44] RJ Zeyen, EL Stromberg, and EL Kuehnast. Long-range aphid transport hypothesis for maize dwarf mosaic virus: history and distribution in Minnesota, USA. Annals of applied biology, 111(2):325–336, 1987.

[45] David RJ Pleydell, Samuel Soubeyrand, Sylvie Dallot, Gérard Labonne, Joël Chadœuf, Emmanuel Jacquot, and Gaël Thébaud. Estimation of the dispersal distances of an aphid-borne virus in and a patchy landscape. PLoS computational biology, 14(4):e1006085, 2018.

[46] M Mcleish, S Sacristan, A Fraile, and F Garcia-Arenal. Scale dependencies and generalism in host use and shape virus prevalence. Proceedings Of The Royal Society B-Biological Sciences, 284(1869), Dec 20 2017.

[47] H Lecoq, F Fabre, B Joannon, C Wipf-Scheibel, C Chandeysson, A Schoeny, and C Desbiez. Search for factors involved in the rapid shift in watermelon mosaic virus (WMV) populations in south-eastern france. Virus Research, 159(2, Si):115–123, Aug 2011.

[48] Cécile Desbiez, Catherine Wipf-Scheibel, Pauline Millot, Karine Berthier, Grégory Girardot, Patrick Gognalons, Judith Hirsch, Benot Moury, Karine Nozeran, Sylvain Piry, et al. Distribution and evolution of the major viruses infecting cucurbitaceous and solanaceous crops in the and French Mediterranean. Virus Research, page 198042, 2020.

[49] C Desbiez, B Joannon, C Wipf-Scheibel, C Chandeysson, and H Lecoq. Recombination in natural populations of watermelon mosaic virus: New agronomic threat or damp squib? Journal Of General Virology, 92(8):1939–1948, Aug 2011.

[50] L Roques, Y Hosono, O Bonnefon, and T Boivin. The effect of competition on the neutral intraspecific diversity and of invasive species. J Math Biol, pages DOI: 10.1007/s00285-014-0825-4, 2014.

## References

[1] Meeker, W. Q., & Escobar, L. A. (1995). Teaching about approximate confidence regions based on maximum likelihood estimation. The American Statistician, 49(1), 48–53.

